# On the rise of AI technologies in virtual screening

**DOI:** 10.64898/2026.01.14.699425

**Authors:** Marco Cecchini, Hryhory Sinenka

## Abstract

AI foundational models for predicting protein-ligand interactions and binding affinities have started to emerge. We challenged Boltz-2 on a difficult dataset constructed on ten ultra-large virtual screening hit lists of pharmacologically relevant targets with *in vitro* binding assays. We show that Boltz-2 is the best classifier, with a success rate twice that of any other rescoring strategy. Ligand classifications by Boltz-2 are straightforward, accurate, efficient and robust, opening to million-compound accurate rankings on commodity resources.

---

Virtual screening (VS) remains a cornerstone of modern drug discovery, enabling the efficient identification of bioactive compounds from vast chemical libraries.^1,2^ Classical virtual-screening approaches, such as structure-based molecular docking, pharmacophore modeling, and ligand-based similarity searches, have long been used to prioritize potential hits for experimental validation. However, these traditional methods often struggle to capture receptor flexibility, solvent effects, and entropic contributions, limiting their predictive power for binding affinity estimations. To address these limitations, rigorous physics-based approaches, mostly based on alchemical free energy simulations, have been developed to predict absolute (ABFE) or relative (RBFE) binding free energies with high accuracy.^3–5^ Despite their proven reliability, these approaches remain computationally demanding and are impractical for large-scale screening.

Recently, the accuracy of non-rigorous methods for rescoring docking hits, i.e., methods that can predict the free energy of binding within ca. 15 minutes of computation, was evaluated on a dataset of ten ultra-large virtual screening hit lists of pharmacologically relevant targets (ULVSH) that underwent subsequent *in vitro* binding assays.^6^ In this context, ligand discrimination between actives and inactives is particularly challenging because: i. virtual hits were selected from docking campaigns with exceptionally high hit rates (i.e., there are no obvious non-binders); ii. active and inactive compounds occupy similar portions of the chemical space and cannot be easily distinguished by chemical similarity; iii. neither actives nor inactives share specific interaction patterns with their target; and iv. the ULVSH dataset is formed of mostly transmembrane proteins, whose modeling and simulation are notoriously challenging.^5^ Interestingly, the work of Sindt *et al*. showed that none of the rescoring functions tested on this dataset could reliably distinguish true binders from experimentally inactive compounds.^6^

Recent advances in artificial intelligence (AI) have begun to transform the drug discovery landscape by offering scalable and accurate alternatives for modeling protein-ligand interactions. AI foundational models for co-folding, e.g., AlphaFold3^7^ or Boltz-2,^8^ can now reliably predict the 3D structure of protein-ligand complexes starting from 2D representations of the protein (FASTA sequence) and the ligand (SMILES format). Moreover, the introduction of an affinity-prediction module in Boltz-2 was shown to achieve accuracy comparable to rigorous physics-based methods, while operating approximately three orders of magnitude faster.^8^ Most recently, using a subset of protein targets from the FEP+ benchmark, Boltz-2 affinity predictions were shown to be on par with ABFE calculations started from corresponding X-ray structures.^9^ These advances provide evidence of a paradigm shift in drug discovery, which opens to scalable and accurate virtual screening workflows alternative to docking with non-rigorous rescoring.

We challenged the Boltz-2 predictive power on the ULVSH dataset. This dataset contains 943 virtual hits (427 true positives, 516 false positives) that were identified in ten ultra-large screening campaigns targeting seven GPCRs, one kinase, one membrane receptor, and one transporter.^6^ Starting from the MOL2 coordinates of the ligands available from Ref.^10^ and the FASTA sequence of the proteins from the PDB, we developed a simple automatic procedure to prepare input files, run Boltz-2 and collect binding affinity predictions for all ligands of the library. By running on a single GPU card of the latest generation (RTX 4500 Ada Generation), Boltz-2 affinity predictions took approximately 100 sec/lig, and the ULVSH library could be processed in 1 day. The Boltz-2 classification performance (i.e., the ability to distinguish true binders from inactive compounds) was assessed by measuring ROC-AUC values per target and comparing with eight popular rescoring strategies including empirical rescoring, machine learning, single-point energy calculations based on polarizable force fields or semi-empirical quantum mechanics, and end-point free-energy simulations from Sindt et al.^6^ To construct ROC curves for Boltz-2, the ligands were ranked based on `affinity_probability_binary` values as recommended for hit identification (see the Boltz-2 documentation^11^); we note that ligand ranking based on the `affinity_pred_value` yields similar results (Table S2).

The results are given in **Table 1**. Remarkably, Boltz-2 predictions outperform all the rescoring strategies tested by Sindt *et al*. ^6^ with an average ROC-AUC of 0.70, which corresponds to a significant increase in classification accuracy. Moreover, if ligand classification is considered successful when the ROC-AUC is higher than 0.7, which is arbitrary but reasonable,^6^ Boltz-2 largely outperforms all other rescoring strategies with a success rate that is more than twice that of the best method in the pool (**Table 1**); note however, that the difference in performances is statistically significant only against docking (Table S4). Last, when repeated on a different machine equipped with a different GPU card (i.e., RTX A4500), Boltz-2 yielded very similar results with absolute ROC-AUC deviations per target below 0.04 units (Table S3) and even slightly better performances (i.e., 83 sec/ligand on RTX A4500 vs 96 sec/ligand on RTX 4500 Ada). Nonetheless, Boltz-2 failed in two out of ten cases (i.e., CNR1 and MTR1A); note that without these two targets, the average ROC-AUC would increase to 0.77.

**Table 1.**
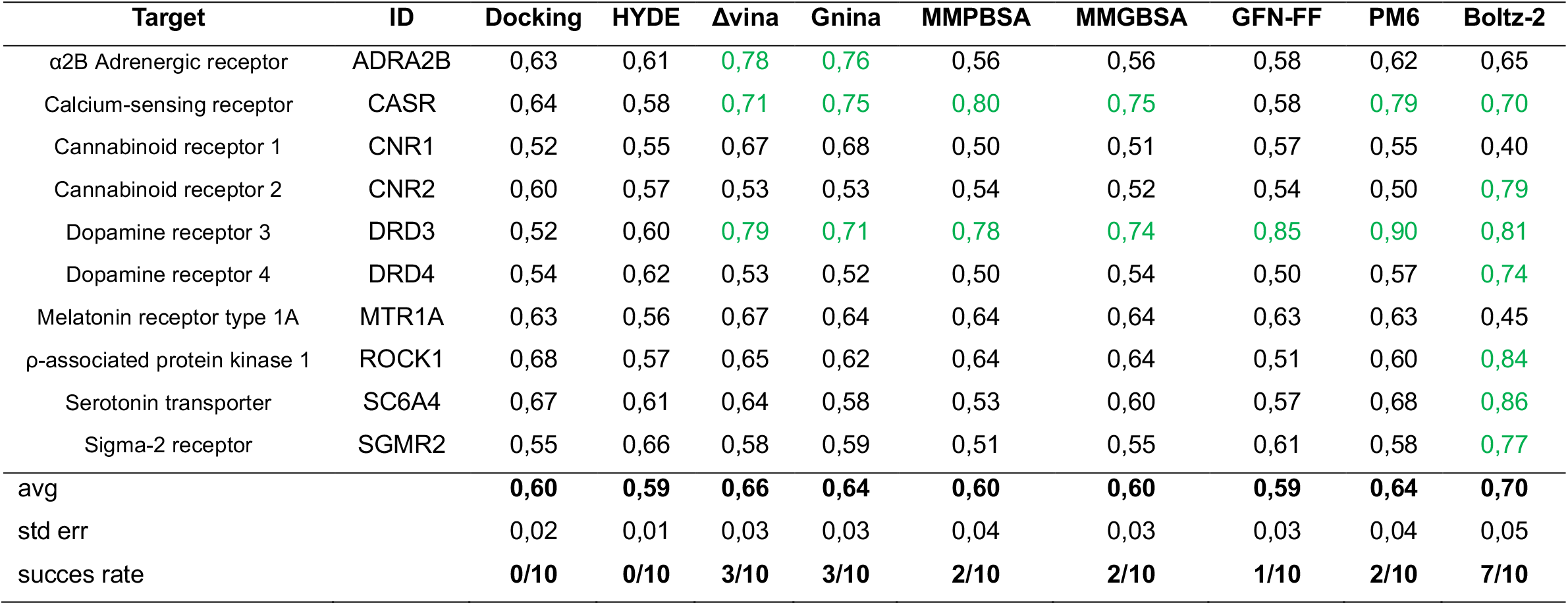
ROC-AUC results for Boltz-2 and a series of commonly used rescoring strategies with the ULVSH dataset. The average ROC-AUC and its standard error over the entire dataset are given at the bottom. Ligand classifications were considered successful when the ROC-AUC was greater than 0.7 (green highlighting). The corresponding ROC curves are shown in Figure S1. ROC-AUC results for all rescoring methods but Boltz-2 were taken from Sindt et al.^6^

To identify potential limitations in our Boltz-2 affinity predictions and possibly correct for them, we re-examined the two unsuccessful targets (CNR1 and MTR1A) and tested different hypotheses. First, since Boltz-2 affinity predictions come with a certain variance in independent calculations, our analysis was repeated 20 times per ligand, and ROC-AUC values were computed using the mean, the best, or the worst prediction to assess Boltz-2 performances in an ensemble framework. Second, because high-resolution structures in complex with ligands were available for all targets (see PDB ID in Ref.^6^), affinity predictions were recomputed by providing Boltz-2 with experimental coordinates as templates (i.e., template conditioning). Third, since most of the targets in ULVSH are multi-domain and the ligand-binding site is always located within one of them, the calculations were repeated by inputting single-domain FASTA sequences corresponding to the 3D coordinates used for docking to remove potential sources of noise. Fourth, to increase the confidence level of the predictions, the analysis was repeated by increasing the quality of the inference parameters (i.e., `recycling_steps=6`, `sampling_steps=1000`, `sampling_steps_affinity=1000`, and `diffusion_samples_affinity=10)` at the price of increased computation. Fifth, under the hypothesis that the true solution could have been missed because of premature convergence, the calculations were repeated by increasing the stochastic character of the search via the `step_scale` and `diffusion_samples` parameters, with the former controlling the diversity of the proposed solutions and the latter setting the number of independent samples used for prediction. The results in Table S5 show that no methodological variant produced a performance improvement relative to the default. One way of affecting the stochastic character of the search (`step_scale`=5 and `diffusion_samples`=20) seems to improve the classification ability for MTR1A (ROC-AUC of 0.66), but it has little to no effect on CNR1. We conclude that Boltz-2 affinity predictions on the ULVSH dataset with default parameters are robust, efficient, and close to optimum. Last, to make sure that the ULVSH compounds were not mis-docked during co-folding, the 3D models predicted by Boltz-2 were compared to the PDB coordinates of existing co-crystals (see PDB codes in Table S1). As shown in Figure S2, all compounds were docked within the crystallographic ligand-binding site. However, when the predicted models were compared to the experimental structures, the ligands appear to be mis-docked (RMSD > 2 Å) in five out of ten cases (Table S6). Surprisingly, we found no correlation between the quality of the structural predictions and the classification performances (Figure S3).

In a second series of experiments, the Boltz-2 performances were assessed in a more drug discovery-like setting. For this purpose, docking hit lists for six of the ten ULVSH targets (i.e., ADRA2B, CNR1, DRD4, MTR1A, SC6A4, and SGMR2) were retrieved from the LSD database^12^ and examined. These lists (ranging from 20 million to 468 million compounds) are substantially larger than those available in ULVSH and provide more representative sets of top-scoring post-docking hits both in terms of chemical diversity and low prevalence of true binders; i.e., the average fraction of active compounds within the top 1,000 is approximately 1% in LSD versus approximately 40% in ULVSH. To keep the calculations tractable, the top 1,000 post-docking hits per target were extracted and mixed with all experimentally confirmed actives, including those that were ranked lower by docking. The resulting hit lists were subsequently re-ranked using the binding probability predicted by Boltz-2. The results show that Boltz-2 rescoring can rescue known binders from low docking ranks, yielding an approximately 4–5-fold enrichment over docking alone (**Table 2**). Moreover, they show that starting at the top-100 fraction, stable enrichments at the different ranks are obtained. These results thus suggest that Boltz-2 rescoring delivers robust enrichments even within the very top fraction of the docking hit list, although the actual enrichment may be target dependent (**Table 2**). When the same analysis was carried out on the ULVSH dataset, the enrichment by Boltz-2 rescoring was still significant but lower; i.e., 1.7-fold over docking at the top 25% (Table S7). Whether the disparity between the ULVSH and LSD results depends on the nature of the classification problem (i.e., rescoring post-docking hits vs rescoring clustered and manually selected compounds) or the design of our experiment with seeded actives from low-docking ranks, remains an open question.

**Table 2.**
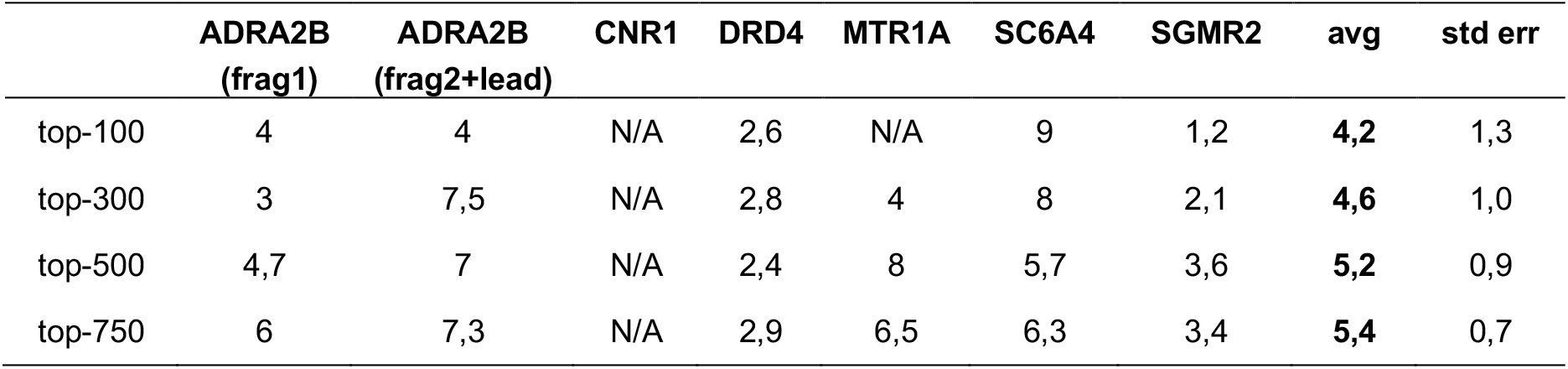
Enrichment by Boltz-2 rescoring of docking hits in the top-ranked fractions of the LSD hit lists used in this work. Boltz-2 rescoring was carried out on the top-1000 docking hits supplemented with all experimentally active compounds available in LSD, including those that were ranked lower by docking. The Boltz-2 enrichments were determined as the ratio between the number of true actives identified by Boltz-2 over that predicted by docking alone. The results (bold font) show that Boltz-2 rescoring yields an approximately four to five fold enrichment over docking alone, even in the top-100 fraction of the hit list.

Binding affinity predictions with Boltz-2 can be generated at a rate of ca. 1,000 ligands per day on a single GPU, corresponding to 20,000 ligands per day, or roughly half a million ligands per month, on a mid-sized cluster with 20 GPUs. Albeit impressive, this throughput is three to four orders of magnitude lower than that of state-of-the-art docking, such as Uni-Dock (0.1 sec/lig),^13^ or machine-learning docking such as KarmaDock (0.01 sec/lig).^14^ Consequently, Boltz-2 remains impractical for exploring the ultra-large chemical spaces that are currently accessible *on demand*^15,16^ and is therefore unlikely to replace docking. However, Boltz-2 calculations performed on 20 GPUs over five days enable the accurate rescoring of approximately 10^5^ compounds with a potentially 2-5-fold enrichment over docking in the top 1% fraction (∼10^3^ compounds). Last, these top-ranked compounds could be further refined using absolute binding free energy calculations (ABFE) initiated from Boltz-2-generated poses, as recently suggested.^9^ Considering accuracy, robustness, and throughput, co-folding methods like Boltz-2 appear to provide an effective rescoring strategy that bridges the gap between ultra-large library screening (∼10^9^ compounds) and lead optimization (∼10^3^ compounds) in a novel and elegant fashion.

During peer review, three relevant contributions on this subject have been released. The first from Bret et al^17^ challenged Boltz-2 on the ULVSH dataset and found consistent with our observations that Boltz-2 is the best ligand classifier among commonly used rescoring strategies. While the same conclusion was reached, this work provides a critical assessment of co-folding methods, highlighting that Boltz-2 affinity predictions are not correlated with pose accuracy, are insensitive to biologically significant mutations in the binding site, and may be exclusively ligand-dependent in extreme scenarios (e.g., when a single alanine was inputted as protein sequence). The second contribution by Shen et al^18^ reported on a systematic assessment of co-folding methods, including AlphaFold3, Boltz-2, and Protenix in virtual screening. Consistent with our observations, it was found that co-folding is superior to conventional docking for ligand classification. However, it was also found that co-folding performances degraded when the assessment was carried out on a new dataset with high-resolution structures released only after the temporal cutoff used for training, when true inactive compounds were incorporated in the test set, or when chemically similar active compounds were excluded from the test set. Last, in the contribution by Kim et al,^19^ ligand classification by co-folding versus docking was assessed on three targets from the LSD dataset using hit lists made of true binders mixed with high-ranking docking inactives. The results show that Boltz-2 did not outperform docking, if not for a somewhat earlier enrichment in the very top fraction of the re-ranking. However, they also show that both methods achieved excellent classification performances within these datasets (i.e., ROC-AUC higher than 0.7), which is unexpected for docking and might not be fully representative of the typical hit-identification scenario; we note that the average ROC-AUC for docking in the ULVSH dataset was 0.6 (**Table 1**). In addition, these and other studies^20^ raise concern about the level of confidence of the co-folding predictions on proteins and ligands that are absent or only marginally present in the original training set. Collectively, recent work by others supports our conclusion that co-folding is superior to conventional docking for ligand classification and may have a role in virtual screening. However, it also raises concern about the domain of applicability of these methods, whose precise definition requires further analysis, particularly in prospective studies.

By redefining the practical limits of virtual screening, the integration of AI-driven technologies like Boltz-2 is challenging the field of drug discovery. For many relevant targets, these methods open to unprecedentedly accurate explorations of large chemical spaces with limited resources and technical requirements. Whether this will lead to a more efficient development of pharmaceuticals is currently unclear, but it is exciting to see how fast the field is evolving and growing.

## Supporting Information

Boltz-2 classification performances using different predictors, runtime parameters or graphics cards; ROC curves for all targets; and structural models of the protein-ligand complexes predicted by Boltz-2 (PDF).

## Data availability

The raw data required to reproduce ligand classification by Boltz-2 in the ULVSH dataset, the structural models by Boltz-2 for all protein-ligand complexes in ULVSH, as well as the different sets of runtime parameters that were used to explore the targets that failed are available at Zenodo. The top 1,000 docking hit lists from the LSD dataset analyzed in this work are also provided [https://zenodo.org/records/19138090].

## Acknowledgments

This work received funding from the European Union’s Horizon 2020 Research and Innovation Program under Marie Sklodowska-Curie Grant Agreement 956314 [ALLODD].

## References

(1) Zhu, H.; Zhang, Y.; Li, W.; Huang, N. A Comprehensive Survey of Prospective Structure-Based Virtual Screening for Early Drug Discovery in the Past Fifteen Years. Int. J. Mol. Sci. 2022, 23 (24), 15961. 10.3390/ijms232415961.

(2) Lyu, J.; Wang, S.; Balius, T. E.; Singh, I.; Levit, A.; Moroz, Y. S.; O’Meara, M. J.; Che, T.; Algaa, E.; Tolmachova, K.; Tolmachev, A. A.; Shoichet, B. K.; Roth, B. L.; Irwin, J. J. Ultra-Large Library Docking for Discovering New Chemotypes. Nature 2019, 566 (7743), 224–229. 10.1038/s41586-019-0917-9.

(3) Fu, H.; Zhou, Y.; Jing, X.; Shao, X.; Cai, W. Meta-Analysis Reveals That Absolute Binding Free-Energy Calculations Approach Chemical Accuracy. J. Med. Chem. 2022, 65 (19), 12970– 12978. 10.1021/acs.jmedchem.2c00796.

(4) Cournia, Z.; Allen, B.; Sherman, W. Relative Binding Free Energy Calculations in Drug Discovery: Recent Advances and Practical Considerations. J. Chem. Inf. Model. 2017, 57 (12), 2911–2937. 10.1021/acs.jcim.7b00564.

(5) Papadourakis, M.; Sinenka, H.; Matricon, P.; Hénin, J.; Brannigan, G.; Pérez-Benito, L.; Pande, V.; Van Vlijmen, H.; De Graaf, C.; Deflorian, F.; Tresadern, G.; Cecchini, M.; Cournia, Z. Alchemical Free Energy Calculations on Membrane-Associated Proteins. J. Chem. Theory Comput. 2023, 19 (21), 7437–7458. 10.1021/acs.jctc.3c00365.

(6) Sindt, F.; Bret, G.; Rognan, D. On the Diiiculty to Rescore Hits from Ultralarge Docking Screens. J. Chem. Inf. Model. 2025, 65 (11), 5553–5566. 10.1021/acs.jcim.5c00730.

(7) Abramson, J.; Adler, J.; Dunger, J.; Evans, R.; Green, T.; Pritzel, A.; Ronneberger, O.; Willmore, L.; Ballard, A. J.; Bambrick, J.; Bodenstein, S. W.; Evans, D. A.; Hung, C.-C.; O’Neill, M.; Reiman, D.; Tunyasuvunakool, K.; Wu, Z.; Žemgulytė, A.; Arvaniti, E.; Beattie, C.; Bertolli, O.; Bridgland, A.; Cherepanov, A.; Congreve, M.; Cowen-Rivers, A. I.; Cowie, A.; Figurnov, M.; Fuchs, F. B.; Gladman, H.; Jain, R.; Khan, Y. A.; Low, C. M. R.; Perlin, K.; Potapenko, A.; Savy, P.; Singh, S.; Stecula, A.; Thillaisundaram, A.; Tong, C.; Yakneen, S.; Zhong, E. D.; Zielinski, M.; Žídek, A.; Bapst, V.; Kohli, P.; Jaderberg, M.; Hassabis, D.; Jumper, J. M. Accurate Structure Prediction of Biomolecular Interactions with AlphaFold 3. Nature 2024, 630 (8016), 493–500. 10.1038/s41586-024-07487-w.

(8) Passaro, S.; Corso, G.; Wohlwend, J.; Reveiz, M.; Thaler, S.; Somnath, V. R.; Getz, N.; Portnoi, T.; Roy, J.; Stark, H.; Kwabi-Addo, D.; Beaini, D.; Jaakkola, T.; Barzilay, R. Boltz-2: Towards Accurate and Eiicient Binding Aiinity Prediction. June 18, 2025. 10.1101/2025.06.14.659707.

(9) Thaler, S.; Wu, Z.; Glass, W. G.; Bradshaw, R. T.; Tossou, P.; Wood, G. P. F. Boltz-ABFE: Free Energy Perturbation without Crystal Structures. arXiv 2025. 10.48550/ARXIV.2508.19385.

(10) The Ultralarge Virtual Screening Hit (ULVSH) Data Set, 2025. http://bioinfo-pharma.u-strasbg.fr/labwebsite/downloads/ULVSH.tgz.

(11) Boltz v2.2.1 Documentation, 2025. https://github.com/jwohlwend/boltz/blob/main/docs/prediction.md.

(12) Hall, B. W.; Tummino, T. A.; Tang, K.; Mailhot, O.; Castanon, M.; Irwin, J. J.; Shoichet, B. K. A Database for Large-Scale Docking and Experimental Results. J. Chem. Inf. Model. 2025, 65 (9), 4458–4467. 10.1021/acs.jcim.5c00394.

(13) Yu, Y.; Cai, C.; Wang, J.; Bo, Z.; Zhu, Z.; Zheng, H. Uni-Dock: GPU-Accelerated Docking Enables Ultralarge Virtual Screening. J. Chem. Theory Comput. 2023, 19 (11), 3336–3345. 10.1021/acs.jctc.2c01145.

(14) Zhang, X.; Zhang, O.; Shen, C.; Qu, W.; Chen, S.; Cao, H.; Kang, Y.; Wang, Z.; Wang, E.; Zhang, J.; Deng, Y.; Liu, F.; Wang, T.; Du, H.; Wang, L.; Pan, P.; Chen, G.; Hsieh, C.-Y.; Hou, T. Eiicient and Accurate Large Library Ligand Docking with KarmaDock. Nat. Comput. Sci. 2023, 3 (9), 789–804. 10.1038/s43588-023-00511-5.

(15) Liu, F.; Wu, C.-G.; Tu, C.-L.; Glenn, I.; Meyerowitz, J.; Kaplan, A. L.; Lyu, J.; Cheng, Z.; Tarkhanova, O. O.; Moroz, Y. S.; Irwin, J. J.; Chang, W.; Shoichet, B. K.; Skiniotis, G. Large Library Docking Identifies Positive Allosteric Modulators of the Calcium-Sensing Receptor. Science 2024, 385 (6715), eado1868. 10.1126/science.ado1868.

(16) Corrêa Veríssimo, G.; Salgado Ferreira, R.; Gonçalves Maltarollo, V. Ultra-Large Virtual Screening: Definition, Recent Advances, and Challenges in Drug Design. Mol. Inform. 2025, 44 (1), e202400305. 10.1002/minf.202400305.

(17) Bret, G.; Sindt, F.; Rognan, D. Assessing Boltz-2 Performance for the Binding Classification of Docking Hits. J. Chem. Inf. Model. 2026, 66 (3), 1511–1521. 10.1021/acs.jcim.5c02630.

(18) Shen, C.; Zhang, X.; Gu, S.; Zhang, O.; Wang, Q.; Du, G.; Zhao, Y.; Jiang, L.; Pan, P.; Kang, Y.; Zhao, Q.; Hsieh, C.-Y.; Hou, T. Unlocking the Application Potential of AlphaFold3-like Approaches in Virtual Screening. Chem. Sci. 2026, 17 (5), 2858–2879. 10.1039/D5SC06481C.

(19) Kim, J.; Correy, G. J.; Hall, B. W.; Rachman, M. M.; Mailhot, O.; Togo, T.; Gonciarz, R. L.; Jaishankar, P.; Neitz, R. J.; Hantz, E. R.; Doruk, Y. U.; Stevens, M. G. V.; Diolaiti, M. E.; Reid, R.; Gopalkrishnan, S.; Krogan, N. J.; Renslo, A. R.; Ashworth, A.; Shoichet, B. K.; Fraser, J. S. Large Scale Prospective Evaluation of Co-Folding across 557 Mac1-Ligand Complexes and Three Virtual Screens. December 28, 2025. 10.64898/2025.12.25.696505.

(20) Škrinjar, P.; Eberhardt, J.; Tauriello, G.; Schwede, T.; Durairaj, J. Have Protein-Ligand Cofolding Methods Moved beyond Memorisation? February 7, 2025. 10.1101/2025.02.03.636309.

