## Supplemental Tables and Figures for "On the rise of AI technologies in virtual screening"

### This document includes:

Tables S1 to S8
Figures S1 to S3

| Full Name | Target ID | Gene name | No. residues | PDB | Uniprot | Organism | First release | No. PDB structures |
|---|---|---|---|---|---|---|---|---|
| α2B Adrenergic receptor | ADRA2B | ADRA2B | 450 | 6K41 | P18089 | *Homo Sapiens* | 04/2020 | 2 |
| Calcium-sensing receptor | CASR | CASR | 1078 | 7M3G | P41180 | *Homo Sapiens* | 03/2021 | 26 |
| Cannabinoid receptor 1 | CNR1 | CNR1 | 472 | 5XR8 | P21554 | *Homo Sapiens* | 11/2016 | 25 |
| Cannabinoid receptor 2 | CNR2 | CNR2 | 360 | 5ZTY | P34972 | *Homo Sapiens* | 01/2019 | 9 |
| Dopamine receptor 3 | DRD3 | DRD3 | 400 | 3PBL | P35462 | *Homo Sapiens* | 11/2010 | 6 |
| Dopamine receptor 4 | DRD4 | DRD4 | 419 | 5WIU | P21917 | *Homo Sapiens* | 10/2017 | 2 |
| Melatonin receptor type 1A | MTR1A | MTNR1A | 350 | 6PS8 | P48039 | *Homo Sapiens* | 04/2019 | 8 |
| Rho-associated protein kinase 1 | ROCK1 | ROCK1 | 1354 | 2ETR | Q13464 | *Homo Sapiens* | 11/2005 | 28 |
| Serotonin transporter | SC6A4 | SLC6A4 | 630 | 6DZZ | P31645 | *Homo Sapiens* | 04/2016 | 29 |
| Sigma-2 receptor | SGMR2 | TMEM97 | 176 | 7MFI | Q3MHW7 | *Bos Taurus* | 12/2021 | 5 |

***Table S1*** *Protein targets from the ULVSH dataset. The date of the first released high-resolution structure and the number of structures currently available from the PDB (March 2026) are given.*

| Target | Docking | Boltz-2 `affinity_pred_value` | Boltz-2 `affinity_probability_binary` |
|---|---|---|---|
| ADRA2B | 0,63 | 0,78 | 0,65 |
| CASR | 0,64 | 0,66 | 0,70 |
| CNR1 | 0,52 | 0,43 | 0,40 |
| CNR2 | 0,60 | 0,69 | 0,79 |
| DRD3 | 0,52 | 0,77 | 0,81 |
| DRD4 | 0,54 | 0,69 | 0,74 |
| MTR1A | 0,63 | 0,48 | 0,45 |
| ROCK1 | 0,68 | 0,77 | 0,84 |
| SC6A4 | 0,67 | 0,83 | 0,86 |
| SGMR2 | 0,55 | 0,72 | 0,77 |
| **avg** | **0,60** | **0,68** | **0,70** |
| **std err** | 0,02 | 0,04 | 0,05 |
| **success rate** | 0/10 | **5/10** | **7/10** |

***Table S2*** *ROC-AUC values for the ULVSH dataset based on Boltz-2 binding affinities or probabilities as recommended for the hit-discovery stage (see documentation at Ref.[11]). The results show that ROC-AUC values are equivalent, albeit binding probabilities provide slightly better classifications. The color code is the same as in Table 1 (Main Text).*

| Target | Docking | Boltz-2 (RTX 4500 Ada) `affinity_probability_binary` | Boltz-2 (RTX A4500) `affinity_probability_binary` |
|---|---|---|---|
| ADRA2B | 0,63 | 0,65 | 0,64 |
| CASR | 0,64 | 0,70 | 0,70 |
| CNR1 | 0,52 | 0,40 | 0,43 |
| CNR2 | 0,60 | 0,79 | 0,77 |
| DRD3 | 0,52 | 0,81 | 0,85 |
| DRD4 | 0,54 | 0,74 | 0,74 |
| MTR1A | 0,63 | 0,45 | 0,49 |
| ROCK1 | 0,68 | 0,84 | 0,84 |
| SC6A4 | 0,67 | 0,86 | 0,83 |
| SGMR2 | 0,55 | 0,77 | 0,76 |
| **avg** | **0,60** | **0,70** | **0,70** |
| **std err** | 0,02 | 0,05 | 0,05 |
| **success rate** | 0/10 | **7/10** | **7/10** |

***Table S3*** *ROC-AUC values for the ULVSH dataset based on Boltz-2 binding probabilities obtained by a different user on a different graphics card. The results show that Boltz-2 predictions with default parameters are robust to the user, the machine, and random errors. The color code is the same as in Table 1 (Main Text).*

| | Docking | HYDE | Δvina | Gnina | MMPBSA | MMGBSA | GFN-FF | PM6 |
|---|---|---|---|---|---|---|---|---|
| p-value | 0,024 | 0,032 | 0,312 | 0,216 | 0,084 | 0,070 | 0,070 | 0,216 |
| significance | yes | yes | no | no | no | no | no | no |

***Table S4*** *Statistical analysis of the Boltz-2 rescoring performance against several rescoring strategies on the ULVSH dataset. For each method, the paired Wilcoxon signed-rank test was run using the ROC-AUC values in Table 1 of the Main Text (n=10). The results show that Boltz-2 rescoring significantly improves over docking and HYDE rescoring, but the difference with the other rescoring strategies is not statistically significant (p > 0.05).*

| Target | Default | Ensemble Mean | Ensemble Best | Ensemble Worst | Template Conditioning | FASTA sequence | High Accuracy | 0.5/5 | 0.5/20 | 5/5 | 5/20 | 5/40 |
|---|---|---|---|---|---|---|---|---|---|---|---|---|
| CNR1 | 0,40 | 0,42 | 0,43 | 0,43 | 0,41 | 0,39 | 0,41 | 0,38 | 0,42 | 0,39 | 0,49 | 0,49 |
| MTR1A | 0,45 | 0,46 | 0,46 | 0,48 | 0,49 | 0,56 | 0,54 | 0,5 | 0,53 | 0,51 | 0,66 | 0,56 |
| **avg** | 0,43 | 0,44 | 0,44 | 0,46 | 0,45 | 0,48 | 0,47 | 0,44 | 0,48 | 0,45 | 0,58 | 0,53 |

***Table S5*** *ROC-AUC values for the two unsuccessful targets of the ULVSH dataset for different setups. Boltz-2 performances were tested by: i. repeating the calculation per ligand 20 times (Ensemble) and considering the mean, the best, and the worst prediction for the ROC curve; ii. using available PDB structures as a template (Template conditioning); iii. using single-domain FASTA sequences; iv. using single-domain FASTA sequences and increasing the quality of the inference parameters (FASTA sequence + High Accuracy); and v. affecting the stochastic character of the search by playing with the temperature of the diffusion and the number of samples used for predictions (`step_scale`/`diffusion_sample`).*

| RMSD [Å] | ADRA2B | CASR | CNR1 | CNR2 | DRD3 | DRD4 | MTR1A | ROCK1 | SC6A4 | SGMR2 |
|---|---|---|---|---|---|---|---|---|---|---|

| | | | | | | | | | | |
|---|---|---|---|---|---|---|---|---|---|---|
| binding site (Cα) | 0,6 | 0,7 | 0,4 | 0,5 | 0,5 | 0,5 | 0,3 | 0,2 | 1,5 | 0,3 |
| ligand binding mode | 2,3 | 2,7 | 0,5 | 0,7 | 2,0 | 2,8 | 0,5 | 0,9 | 3,1 | 5,8 |

***Table S6***. *Structural analysis of the Boltz-2 co-folding predictions for ULVSH based on existing co-crystals. The PDB codes of the structures used for this analysis are given in Table S1. The RMSD of the ligand binding site and the ligand binding mode from the X-ray coordinates are given upon optimal superimposition of the Cα atoms of the protein within 10 Å from the crystallographic ligand-binding pose. The numbers in red highlight ligand mis-docking (RMSD > 2 Å).*

| ULVSH | ADRA2B | CASR | CNR1 | CNR2 | DRD3 | DRD4 | MTR1A | ROCK1 | SC6A4 | SGMR2 | avg | std err |
|---|---|---|---|---|---|---|---|---|---|---|---|---|
| top-25% | 1,2 | 1,1 | 0,8 | 3,1 | 2,5 | 1,3 | 1,0 | 2,6 | 1,8 | 1,2 | 1,7 | 0,2 |
| top-50% | 1,0 | 1,3 | 0,8 | 1,7 | 1,5 | 1,2 | 0,8 | 1,5 | 1,4 | 1,0 | 1,2 | 0,1 |
| top-75% | 1,1 | 1,1 | 0,8 | 1,4 | 0,9 | 1,2 | 0,8 | 1,2 | 1,3 | 1,1 | 1,1 | 0,1 |

***Table S7*** *Relative enrichment by Boltz-2 rescoring over docking in the ULVSH dataset. Enrichment factors were determined as the ratio between the number of true actives identified by Boltz-2 over that predicted by docking at various fractions of the hit lists (i.e. 25%, 50%, and 75%). The results (in red) show that Boltz-2 rescoring yields an average enrichment of 1.7 at top-25%.*

| | N of actives | top-1k | top-2k | top-5k | top-10k | top-20k | top-50k | top-100k |
|---|---|---|---|---|---|---|---|---|
| ADRA2B (frag1) | 21 | 4 | 9 | 13 | 14 | 15 | 17 | 18 |
| ADRA2B (frag2+lead) | 29 | 3 | 4 | 10 | 15 | 21 | 23 | 24 |
| CNR1 | 16 | 0 | 0 | 1 | 2 | 4 | 13 | 13 |
| DRD4 | 80 | 26 | 33 | 54 | 54 | 54 | 62 | 68 |
| MTRA1 | 14 | 2 | 3 | 8 | 9 | 13 | 14 | 14 |
| SC6A4 | 20 | 5 | 7 | 11 | 17 | 20 | 20 | 20 |
| SGMR2 | 201 | 46 | 47 | 63 | 73 | 100 | 121 | 121 |

***Table S8*** *Hit rates at different fractions of the docking hit lists from LSD considered in this study. For the α2B adrenergic receptor (ADRA2B), two of the three docking hit-lists available from LSD (i.e., frag2 and lead) were merged as their screening was done under the same docking settings, unlike frag1.*

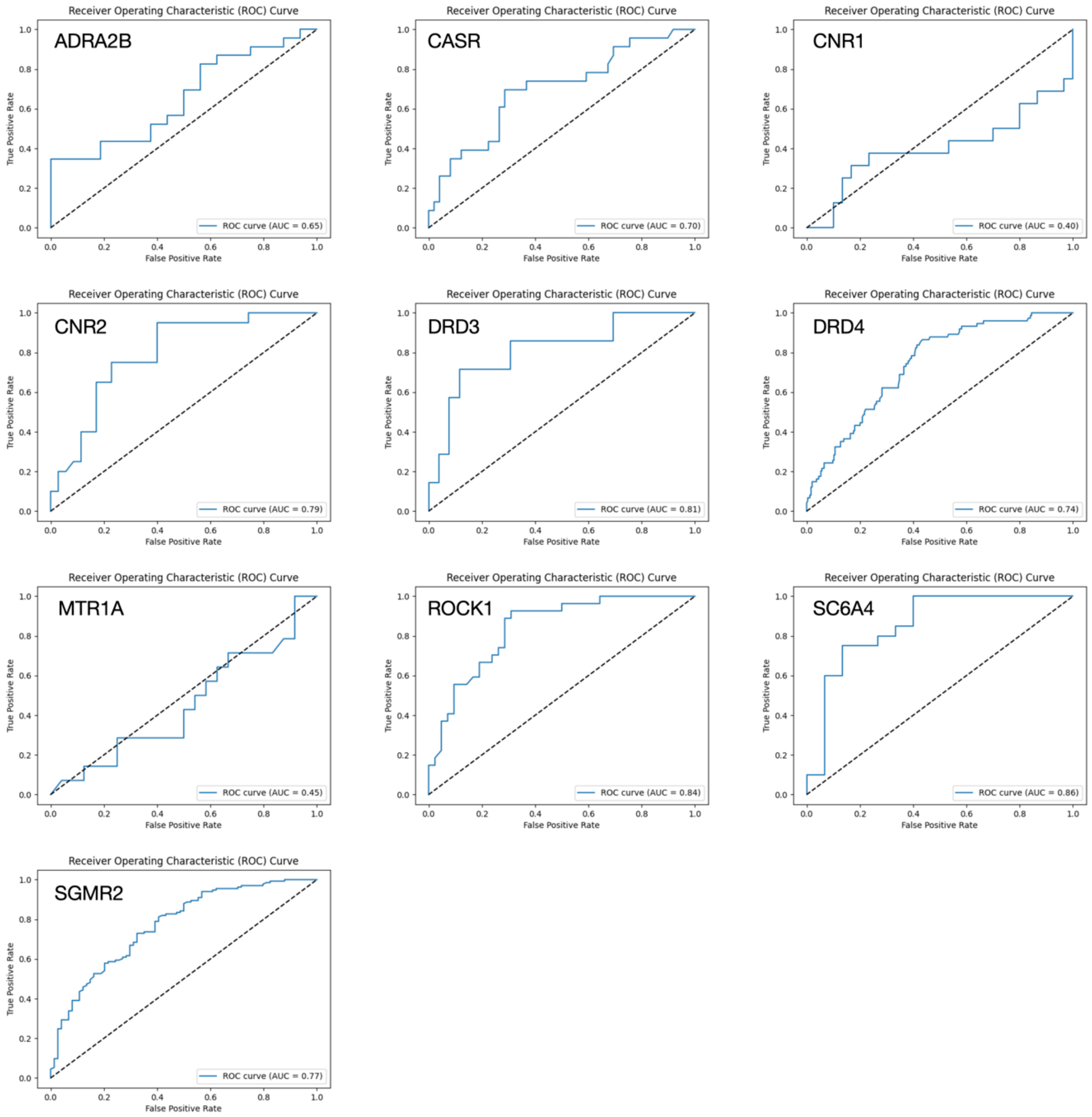


***Figure S1*** *ROC curves for all targets of the ULVSH dataset based on Boltz-2 binding probabilities. The corresponding ROC-AUC values are given in Table 1 of the Main Text.*

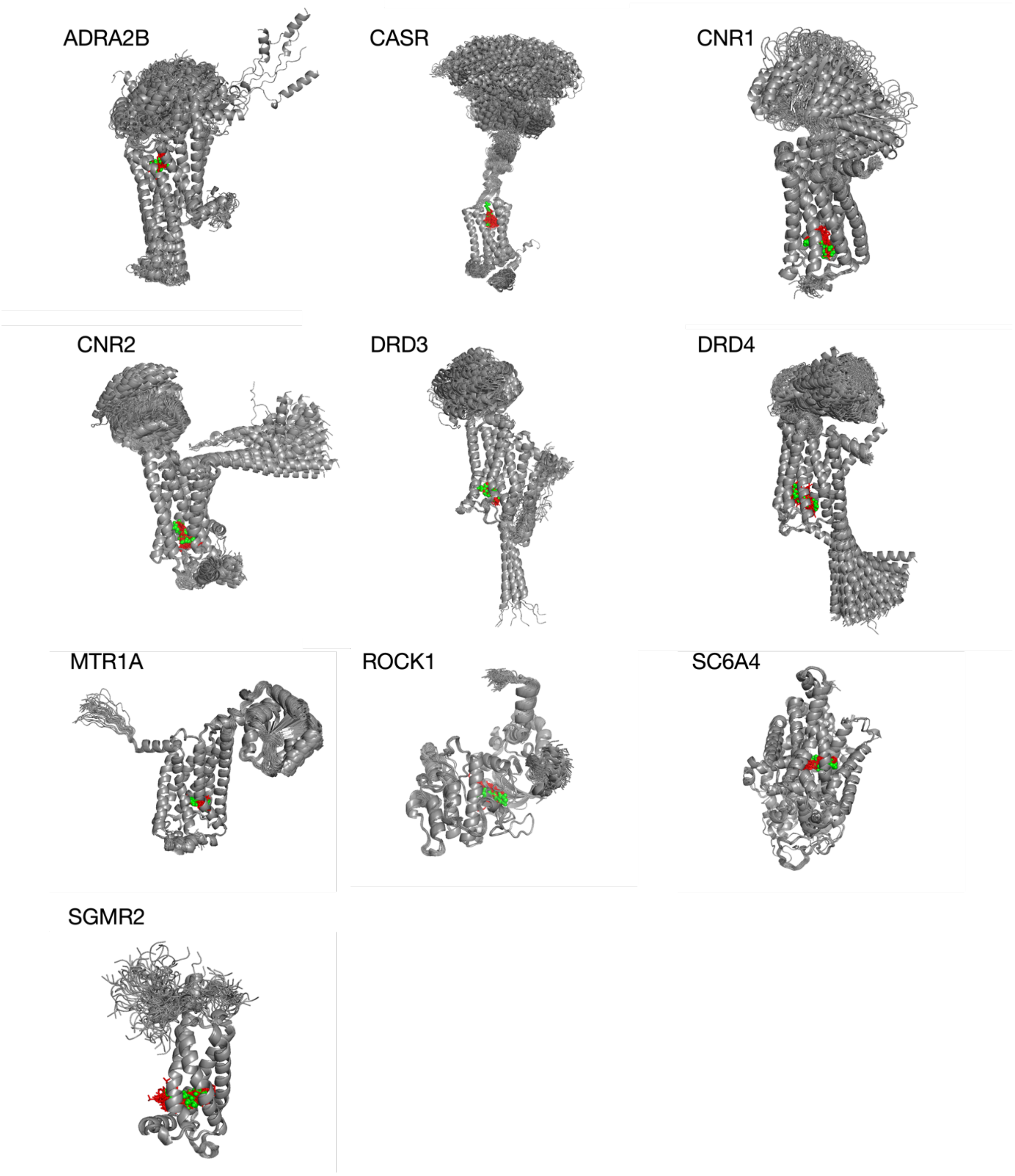


***Figure S2*** *Structural models predicted by Boltz-2 and used for the binding affinity predictions. The co-folded ligands are shown as red sticks. The crystallographic ligand from the PDB is shown as green spheres. The structures show that all co-folded ligands were docked to the crystallographic ligand-binding site. The alignment was done by fitting the C-alpha coordinates of the Boltz-2 co-folding models to the crystallographic binding site, i.e. all C-alpha atoms within 10 Å of the X-ray ligand.*

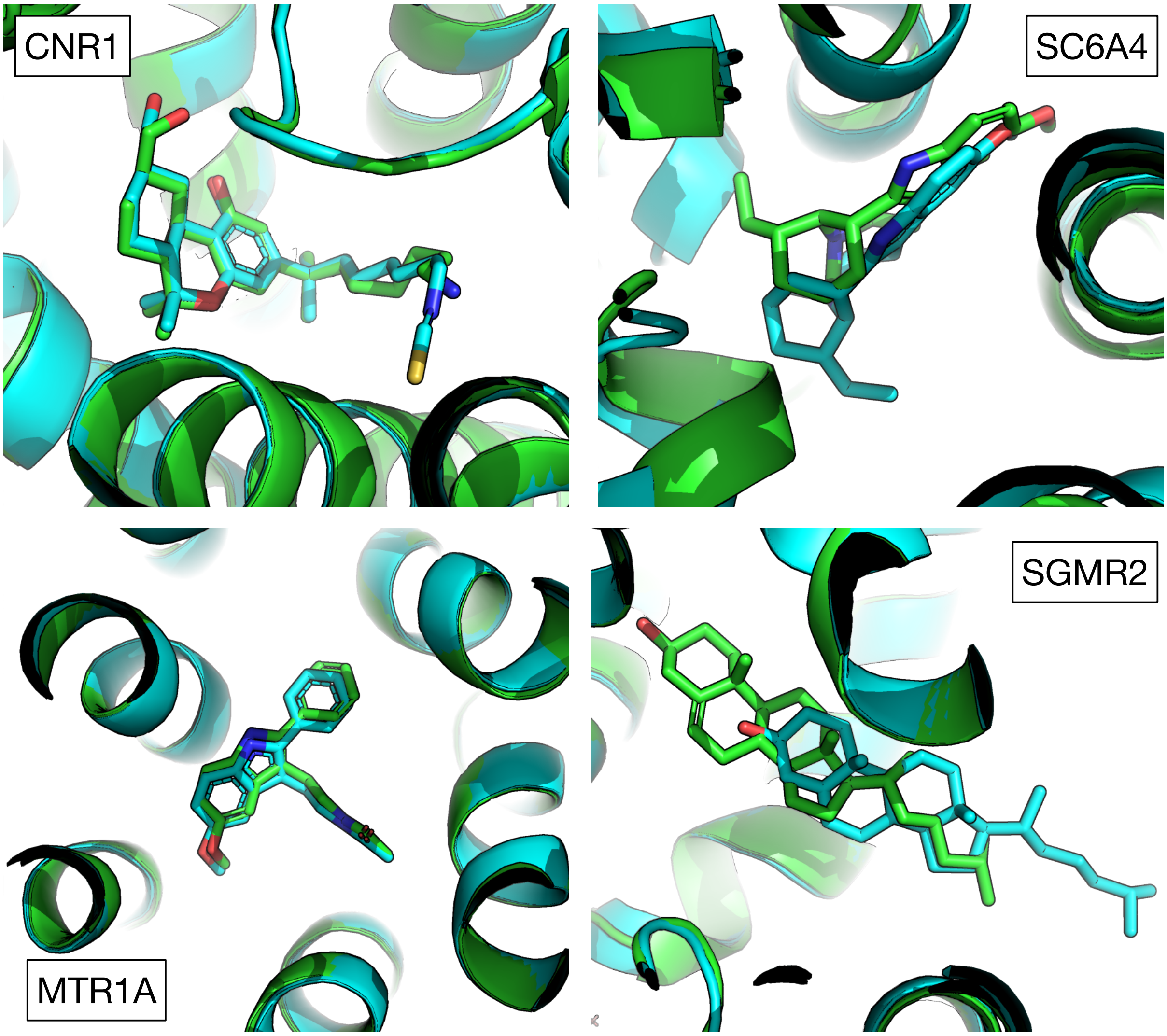


***Figure S3*** *Structural analysis of the Boltz-2 co-folding predictions for existing co-crystals for four targets in the ULVSH dataset. Boltz-2 results for MTR1A and CNR1 are shown on the left. Those for SC6A4 and SGMR2 are shown on the right. This analysis shows that the predicted binding mode (cyan) of the crystallographic ligands in two targets that failed (i.e., CNR1 and MTR1A) perfectly match those in the PDB. In sharp contrast, those predicted in two targets that did not fail (i.e., SC6A4 and SGMR2) are largely mis-docked.*
